# An open-source control system for in vivo fluorescence measurements from deep-brain structures

**DOI:** 10.1101/399329

**Authors:** Scott F. Owen, Anatol C. Kreitzer

## Abstract

**Background:** Intracranial photometry through chronically implanted optical fibers is a widely adopted technique for measuring signals from fluorescent probes in deep-brain structures. The recent proliferation of bright, photo-stable, and specific genetically-encoded fluorescent reporters for calcium and for other neuromodulators has greatly increased the utility and popularity of this technique.

**New Method:** Here we describe an open-source, cost-effective, microcontroller-based solution for controlling optical components in an intracranial photometry system and processing the resulting signal.

**Results:** We show proof-of-principle that this system supports high quality intracranial photometry recordings from dorsal striatum in freely moving mice. A single system supports simultaneous fluorescence measurements in two independent color channels, but multiple systems can be integrated together if additional fluorescence channels are required. This system is designed to work in combination with either commercially available or custom-built optical components. Parts can be purchased for less than one tenth the cost of commercially available alternatives and complete assembly takes less than one day for an inexperienced user.

**Comparison with Existing Method(s):** Currently available hardware draws on a variety of commercial, custom-built, or hybrid elements for both optical and electronic components. Many of these hardware systems are either specialized and inflexible, or over-engineered and expensive.

**Conclusions:** This open-source system increases experimental flexibility while reducing cost relative to current commercially available components. All software and firmware are open-source and customizable, affording a degree of experimental flexibility that is not available in current commercial systems.

## INTRODUCTION

Intracranial volumetric imaging of population-level fluorescence signals through optical fibers (photometry) has been possible for more than a decade (Adelsberger, Garaschuk et al. 2005). Recently, however, the utility and popularity of this technique has increased dramatically, through improvements in brightness and signal-to-noise of genetically-encoded fluorescence indicators (Cui, Jun et al. 2013, Gunaydin, Grosenick et al. 2014, Lerner, Shilyansky et al. 2015, Kim, Yang et al. 2016), most notably the GCaMP family of genetically-encoded calcium indicators (Akerboom, Chen et al. 2012, Chen, Wardill et al. 2013). In addition, the growing availability of specific genetically encoded fluorescent indicators for detection of neuromodulators (Jing, Zhang et al. 2018, Patriarchi, Cho et al. 2018, Sun, Zeng et al. 2018), voltage (Gong, Huang et al. 2015), and complementary cell signaling processes (Miesenbock, De Angelis et al. 1998, Okumoto, Looger et al. 2005, Li and Tsien 2012, Gong, Wagner et al. 2014, San Martin, Ceballo et al. 2014, Marshall, Li et al. 2016) highlights the rapidly expanding versatility of this technique. These advances have driven the development of commercially available hardware capable of replacing custom-built systems and are lowering the barriers to entry for new labs to adopt this technique. However, the electronics to control these commercially available optical systems are either over-engineered (capable of acquiring many more channels at much higher acquisition rates than required) or lack the flexibility to be customized for all applications. Open-source solutions can address these shortcomings by providing a complete, cost-effective system that can be customized as necessary by each user for the needs of any specific experiment.

A typical photometry system consists of three components: (1) a short length of optical fiber that is stereotaxically guided to the brain region of interest and then permanently cemented to the skull, (2) a set of optical components to generate excitation light and detect fluorescence emission, and (3) electronics to control light delivery and digitize the resulting fluorescence signal (Figure 1). For the first component, the most widely used implants are essentially identical to the fiber-ferrule assemblies used for light delivery in optogenetics experiments, which have been established and iteratively improved by a large community for more than a decade (Aravanis, Wang et al. 2007, Sparta, Stamatakis et al. 2011). For the second component, recently-released optical elements that are specialized for this type of recording offer excellent ease-of-use and signal-to-noise at a cost that is comparable to more cumbersome first-generation setups (Gunaydin, Grosenick et al. 2014, Simone, Fuzesi et al.2018). For the third component, a specialized set of electronics are required to control the fluorescence excitation light and to process the resulting fluorescence emission signal. Here we describe the design, construction, and implementation of a low-cost, open-source system for control, recording, on-line visualization, and post-hoc analysis of *in vivo* photometry experiments. This system is designed to operate together with commercially available optical components (e.g. Doric Lenses, Thorlabs) as well as open-source (Delmans 2018, Simone, Fuzesi et al. 2018) and hybrid systems (https://sites.google.com/view/multifp/hardware), and has the ability to monitor synchronization signals to support integration of recordings with additional stimulation or recording equipment. This design supports two independent color channels and can be assembled by an inexperienced user in less than a day, for approximately one tenth the cost of commercially available systems. Most importantly, as both physiological and behavioral experiments become increasingly complex, the open-source architecture allows unlimited range to expand, modify, customize and adapt the system to suit the requirements of any specific experiment.

**Figure 1.**
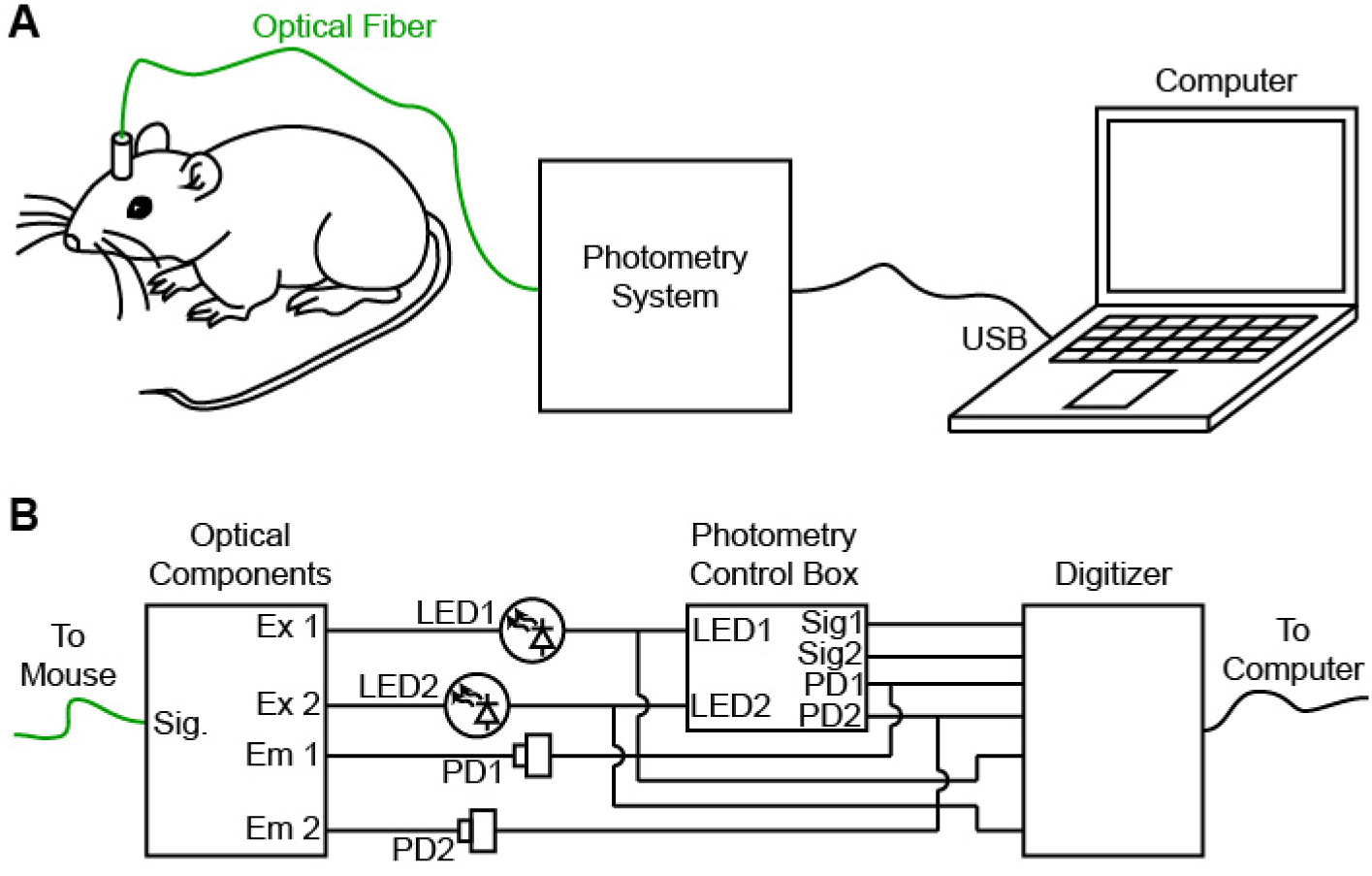
Design of a low-cost open-source photometry control system. A, Schematic of experimental design in which the photometry system delivers fluorescence excitation light, measures fluorescence emission and passes a digitized signal to the computer. B, Individual components of the photometry system include optical components (left, dichroic mirrors, fiber couplers), photometry control box (center) and digitizer for recording (right).

## RESULTS

A key challenge in the design and implementation of an intracranial photometry fluorescence recording system is establishing the extent to which the measurement reflects the true fluorescence signal in the brain. Two distinct approaches have been established to address this problem. The first approach splits the fluorescence signal into multiple different channels based on wavelength and relies on spectral unmixing to isolate the true fluorescence signal in one expected channel from non-specific background fluctuations that are expected to spread across multiple wavelengths (Cui, Jun et al. 2013, Cui, Jun et al. 2014, Meng, Zhou et al. 2018). An alternative approach, which avoids the requirement for complex time-correlated single-photon counting (TCSPC), is to rapidly oscillate the excitation light and use post-hoc processing to isolate the specified component of the fluorescence signal (Gunaydin, Grosenick et al. 2014). In this second approach, it is the temporal characteristics of the fluorescent signal (frequency and phase of the amplitude oscillation) rather than the wavelength that are used to isolate the fluorescence emission signal from background light. Both methods are effective for removing background light and noise sources from the fluorescence signal, or for isolating distinct color channels from one another within the same preparation. The system described here employs the second approach, oscillating the fluorescence excitation light independently for two separate color channels. Notably, neither approach is completely immune to noise sources or experimental confounds, especially those caused by movement of the animal or flexing of the fiber-optic cable. To identify and exclude those sources of noise, it is important to record from a stable, non-fluctuating fluorescence source such as EGFP or tdTomato. These control recordings can be performed in a separate control cohort of animals, or alongside the primary recordings using a second color channel as described below.

The electronics required to operate a photometry system require little more than a sine-wave generator and a low-cost digitizer. However, it is useful in the day-to-day execution of experiments to have a dynamic, on-line estimate of the fluorescence signal amplitude. Here, we describe a newly developed, open-source system that supports all these features. This system is based on commercially available microcontrollers and electronic components, freely-available open-source firmware (for the MBED microcontroller) and post-hoc analysis software (Matlab script). It can be assembled in less than a day by an inexperienced user for a total cost of $500-$1,000; about one tenth the cost of current commercially available systems.

The core of this system is an MBED Cortex LPC1768 programmable microcontroller (Figure 2A). This microcontroller generates two continuously oscillating sine wave outputs to drive excitation light for two independent fluorescence channels (LED). The amplitude (0-3.3 V) and frequency (0-500 Hz) of each oscillation is controlled through four user-defined command voltages (variable resistors). Digital displays provide a readout of each oscillation frequency so that harmonic interference between fluorescence excitation and oscillating signals in the environment (e.g. 60 Hz room lights, or optogenetic stimulation pulse trains) can be avoided.

**Figure 2.**
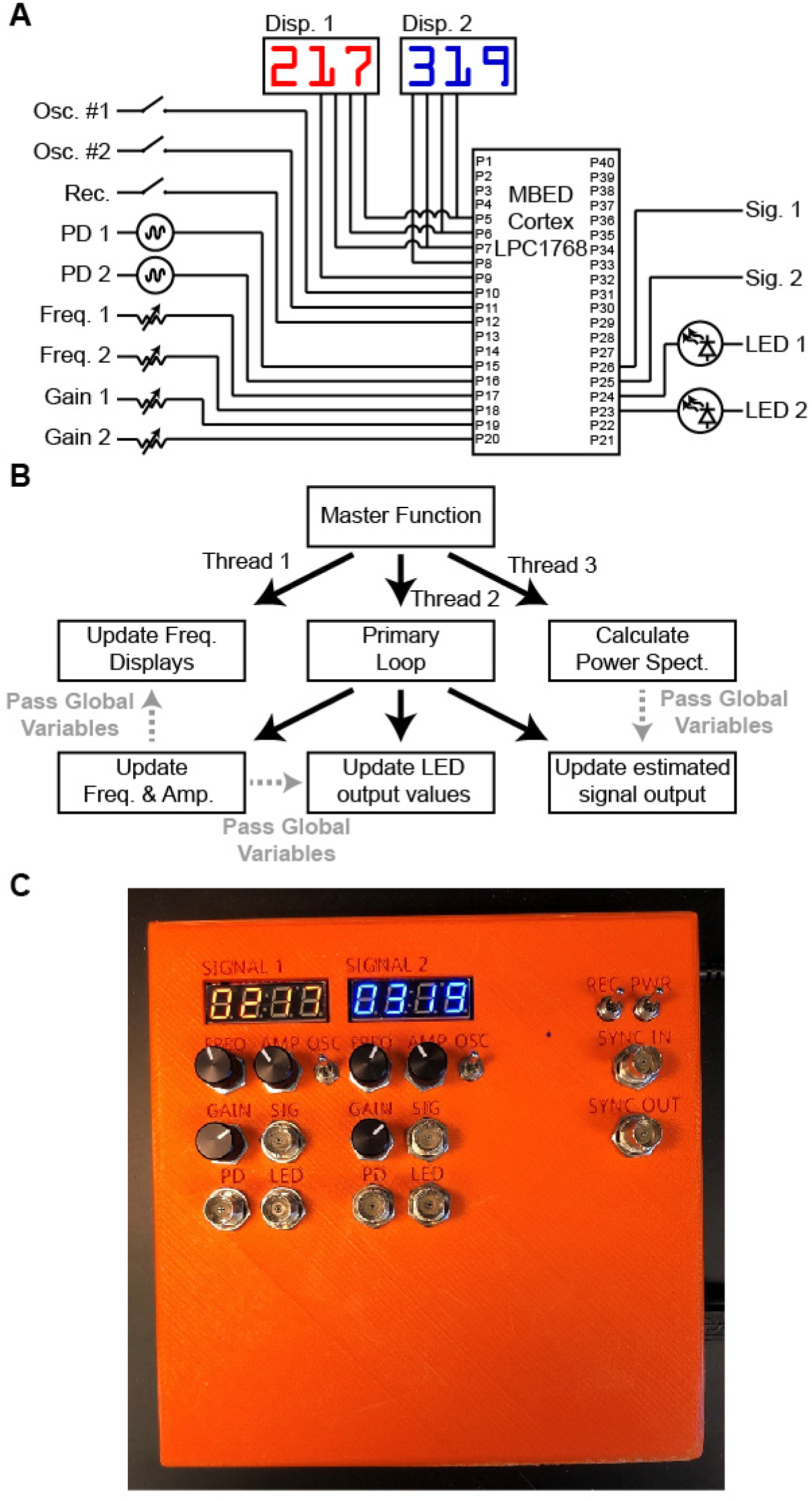
Microcontroller firmware for LED control and on-line readout of fluorescence signal. A, Connectivity of MBED Cortex LPC1768 microcontroller. B, Schematic of microcontroller firmware. C, Constructed photometry control box.

Two switches pause the LED oscillations for each channel. This allows excitation light power to be measured accurately during the setup of an experiment and permits the user to perform experiments with steady rather than oscillating excitation light if desired. A “recording” switch suspends all user input to ensure stable conditions during recording or execution of an experiment, even if the user bumps a knob. The microcontroller also monitors the oscillating fluorescence signal from each channel and calculates a Fast-Fourier Transform (FFT) from each channel to determine the signal power at the frequency of the excitation LED oscillation. This provides a continuously updated estimate of this fluorescence signal for each channel. On- line estimates of the fluorescence signal are calculated by the microcontroller approximately once every ˜80 ms, using a ˜100 ms sliding window. The time delay caused by the filter lag and the microcontroller processing steps introduce a total lag of ˜250 ms for the estimated signal relative to the “true” fluorescence calculated post-hoc using a zero-lag filter. This delay is relatively small compared to the time course of most bulk fluorescent transients recorded *in vivo* (Cui, Jun et al. 2013, Gunaydin, Grosenick et al. 2014, Lerner, Shilyansky et al. 2015, Sun, Zeng et al. 2018), but is important to correct with appropriate post-hoc processing in cases where fine timescale alignment of fluorescent transients to behavior or to complementary physiological signals is required.

Simultaneous execution of these operations on a single microcontroller is made possible by the multi-threaded structure supported on the MBED cortex LPC1768 (Figure 2B). Briefly, three continuously repeating loops run in parallel. The first loop updates the frequency displays. The second loop calculates the amplitude of the oscillating fluorescent signal. The third loop controls the sine wave generation for the oscillating excitation LEDs.

All essential signals are recorded and digitized at ˜5 kHz with a commercially available digitizer (National Instruments USB-6009) and freely available software (WinEDR, http://spider.science.strath.ac.uk/sipbs/software_ses.htm), including the excitation LED driver signals, the raw fluorescent emission signal, and the on-line estimates of the fluorescent signals. Simultaneous digitization of additional synchronization inputs allows the fluorescence signal to be aligned precisely to other physiological manipulations or behavior. The entire system, including digitizer, is contained in a compact box and can be readily moved from one experimental setup to another as needed (Figure 2C).

Assembly of this system is rapid and straightforward, requiring less than a day for a naïve user with modest soldering experience. Briefly, the microcontroller, BNC connectors, switches, variable resistors, resistors, capacitors, and connectors are soldered into the custom-designed printed circuit board. The MBED microcontroller is then connected to a computer via USB (it appears as a USB “thumb drive”) and the firmware is and copied onto the microcontroller. The driver for the digitizer card is installed on the computer as well as the acquisition software (WinEDR, link above). The digitizer is plugged into the printed circuit board and connected to the computer by USB. The control system is then placed into its enclosure. Two BNC cables connect the control system to the LED drivers, and a further two BNC cables connect the photodiodes to the control system. A USB cable connects the digitizer to the acquisition computer, and a 5V power supply provides power. A complete list of parts, pre-compiled firmware, source-code, and all materials as well as a more detailed description of the assembly process is available here https://hackaday.io/project/160397).

To test this system, we injected adeno-associated virus to express the genetically-encoded calcium indicator GCaMP6m in a Cre-dependent manner (AAV-Flex-GCaMP6m) into the dorso-medial striatum of an A2a-Cre BAC transgenic mouse. This drives expression of the calcium indicator selectively in striatal medium spiny neurons (MSNs) belonging to the indirect pathway (Cui, Jun et al. 2013). We implanted a short length of optical fiber (400 µm diameter) approximately 100 µm above the center of the virus infection zone and cemented the implant to the mouse skull. Fluorescence excitation light was generated by a pair of LEDs (Doric lenses) and routed through a specialized “Fluorescence mini cube” (Doric Lenses) containing pre-aligned lenses, fiber couplers, and dichroic mirrors to split excitation and emission channels based on wavelength. Emission light was collected separately for each fluorescence channel with a pair of Newport 2151 Femtowatt detectors.

One standard experimental configuration requiring a two-color photometry system uses a green channel to track a fluorescent sensor (e.g. GCaMP) and a red channel to simultaneously track a static control indicator (e.g. tdTomato). In the specific case of GCaMP indicators, however, an elegantly simple alternative control is possible. Excitation of GCaMP at the standard wavelength for green fluorescence (465 nm) produces a robust calcium-dependent signal; in contrast, blue-shifted excitation light (405 nm) drives a calcium-independent (isosbetic) fluorescence from the same indicator. This second isosbestic wavelength provides a calcium-independent signal to control for preparation stability.

We therefore delivered excitation light at two separate wavelengths: 405 nm to drive isosbestic fluorescence (Figure 3, left column), and 465 nm to drive a calcium-dependent fluorescent signal (Figure 3, right column). To ensure adequate separation of the two fluorescent signals, the intensity of light at each wavelength was independently modulated at different frequencies. Importantly, each oscillation frequency must be rapid relative to the timescale of the signal fluctuations to be detected, but slower than the detection speed of the photodiode and digitizer. In our hands, oscillation frequencies in the range of 100-400 Hz meet these criteria. To avoid generating harmonic interference, each frequency must not be a multiple (or near multiple) of the other frequency, or of any oscillating signal in the environment, such as room lights (60 Hz) or optogenetic stimulation pulse trains. Based on these criteria and empirical testing, we selected 217 Hz as the oscillation frequency for the 465 nm calcium-dependent fluorescence channel, and 319 Hz as the frequency for the 405 nm isosbestic signal.

**Figure 3.**
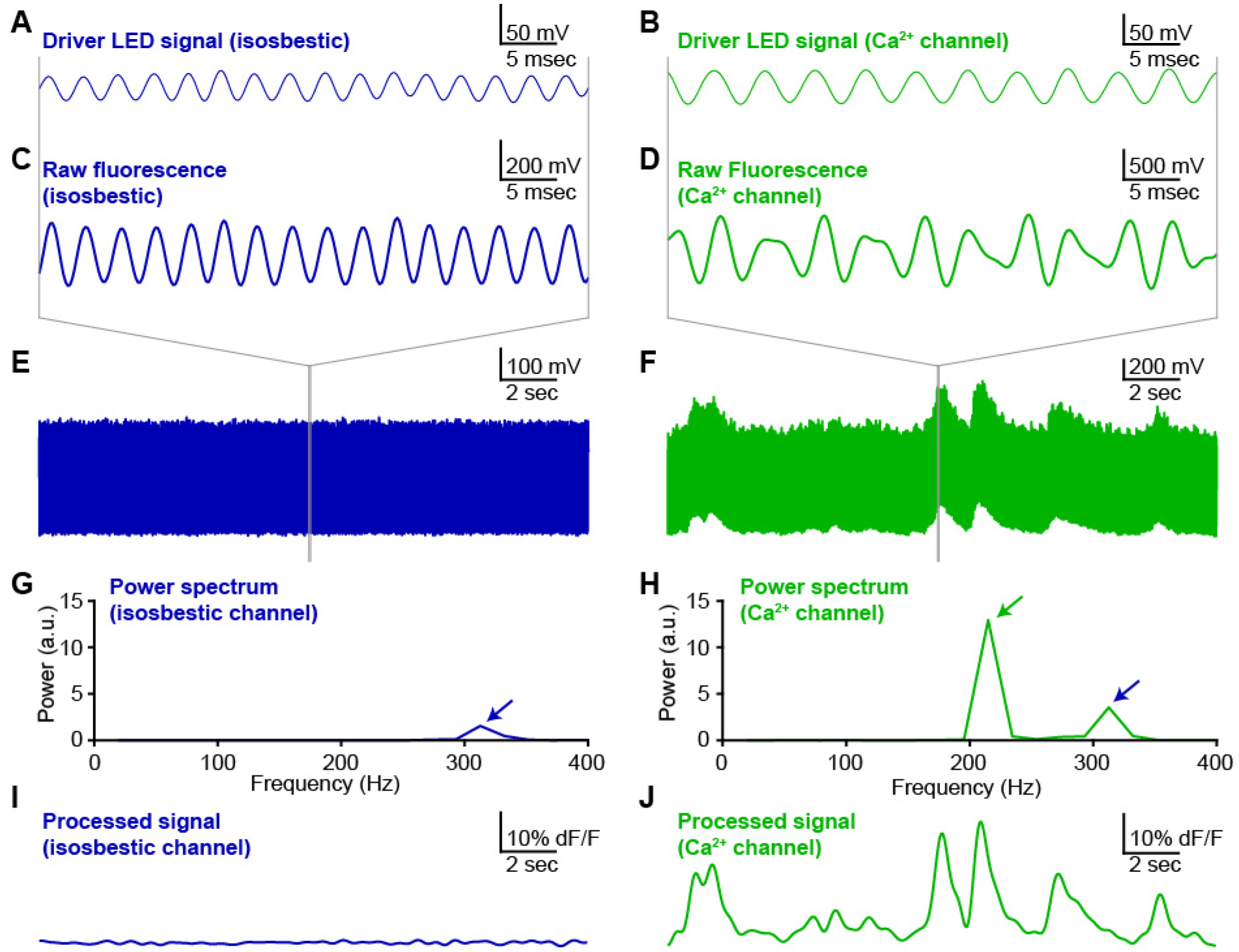
Oscillating excitation light minimizes background fluorescence and channel cross-talk. Exemplar photometry recording and analysis from dorsomedial striatum of A2A-cre mouse expressing GCaMP6m in medium spiny neurons (MSNs) of the indirect pathway. A,B, Oscillating driver signals for isosbestic channel (Panel A, 405 nm; oscillating at 319 Hz) and calcium-sensitive channel (Panel B, 465 nm; oscillating at 217 Hz). C,D, Fluorescence signal from isosbestic channel (Panel C) and calcium-sensitive channel (Panel D) at expanded time scale to show phase-locked oscillations in fluorescence emission relative to each LED driver signal (panels A,B above). E,F, Fluorescence measurements for each channel at long time scale to show stability of isosbestic channel (Panel E) relative to calcium-sensitive fluctuations in fluorescence (Panel F). G,H, Power spectra calculated from a discrete 50 ms window of each fluorescence emission signal. Blue arrows indicate the peak corresponding to the isosbestic channel (405 nm excitation light; oscillation at 319 Hz). Green arrow indicates peak corresponding to calcium-dependent channel (465 nm excitation light; oscillation at 217 Hz). I,J, Fluorescence signal for each channel calculated by post-hoc processing over a sliding window (see methods).

In our validation recording, the fluorescence emission from the isosbestic channel (405 nm excitation light) appeared as an oscillation that was phase-locked to the LED driver signal (Figure 3A,C,E). The fluorescence emission signal from the calcium-sensitive channel (465 nm excitation light) exhibited a more complex structure (Figure 3B,D,F), consistent with this signal arising from a summation of the isosbestic and calcium-sensitive fluorescence signals. The cross-talk between these two channels was readily isolated by using a Fast-Fourier Transform (FFT) to calculate the local power spectrum of the signal across a sliding time window. The single peak in the power spectrum for the isosbestic signal confirmed the specificity of this fluorescence measurement (Figure 3G). In contrast, the power spectrum from the calcium-sensitive fluorescence channel contained two fully separated peaks corresponding to the calcium-sensitive fluorescence (green arrow) and the cross-talk from the isosbestic fluorescence signal (blue arrow) (Figure 3H).

The amplitude of the calcium-sensitive fluorescence signal was calculated by measuring peak of the power spectrum at the frequency dictated by the oscillation of the calcium-sensitive fluorescence excitation light (green arrow). By repeating this calculation over a sliding time window, the fluorescence signal for each channel was calculated as a function of time, independent of the other fluorescence channel or background light contamination. As expected, the fluorescence signal from the isosbestic channel was stable over time (Figure 3I), while the calcium-sensitive fluorescence channel showed robust fluctuations (Figure 3J). The size, shape and time-course of these fluctuations was consistent with previously reported calcium transients in similar preparations (Cui, Jun et al. 2013, Meng, Zhou et al. 2018). In some cases, such as when significant fluctuations are detected in the control fluorescence channel, it may be desirable to normalize the calcium-dependent fluorescence by the instantaneous control signal amplitude to correct for this noise in the primary signal. When the control signal is stable, however, it often makes sense to skip this normalization, because dividing one experimentally measured signal by another will increase the noise in the final signal.

To align calcium fluorescence signals to behavior, we recorded fluorescence continuously for 10 min (Figure 4A,B) while monitoring animal locomotion using an overhead camera (Figure 4C). Synchronization pulses generated by the behavioral monitoring system (Ethovision 10, Noldus Inc) were digitized by the photometry recording system together with the fluorescence measurements (see methods). Consistent with previous reports (Cui, Jun et al. 2013, Meng, Zhou et al. 2018), turns or orienting movements towards the direction contralateral to the recording site in dorsomedial striatum were correlated with peaks in the fluorescent signal on the calcium-dependent channel (Figure 4D). In contrast, turns or orienting movements towards the ipsilateral side were associated with troughs, or the absence of peaks, in the fluorescence signal (Figure 4E). No peaks or behaviorally-aligned changes were detected in the isosbestic control signal, consistent with the transients in the calcium-dependent channel representing well-isolated calcium signals with minimal contamination from movement artifacts or environmental light sources.

**Figure 4.**
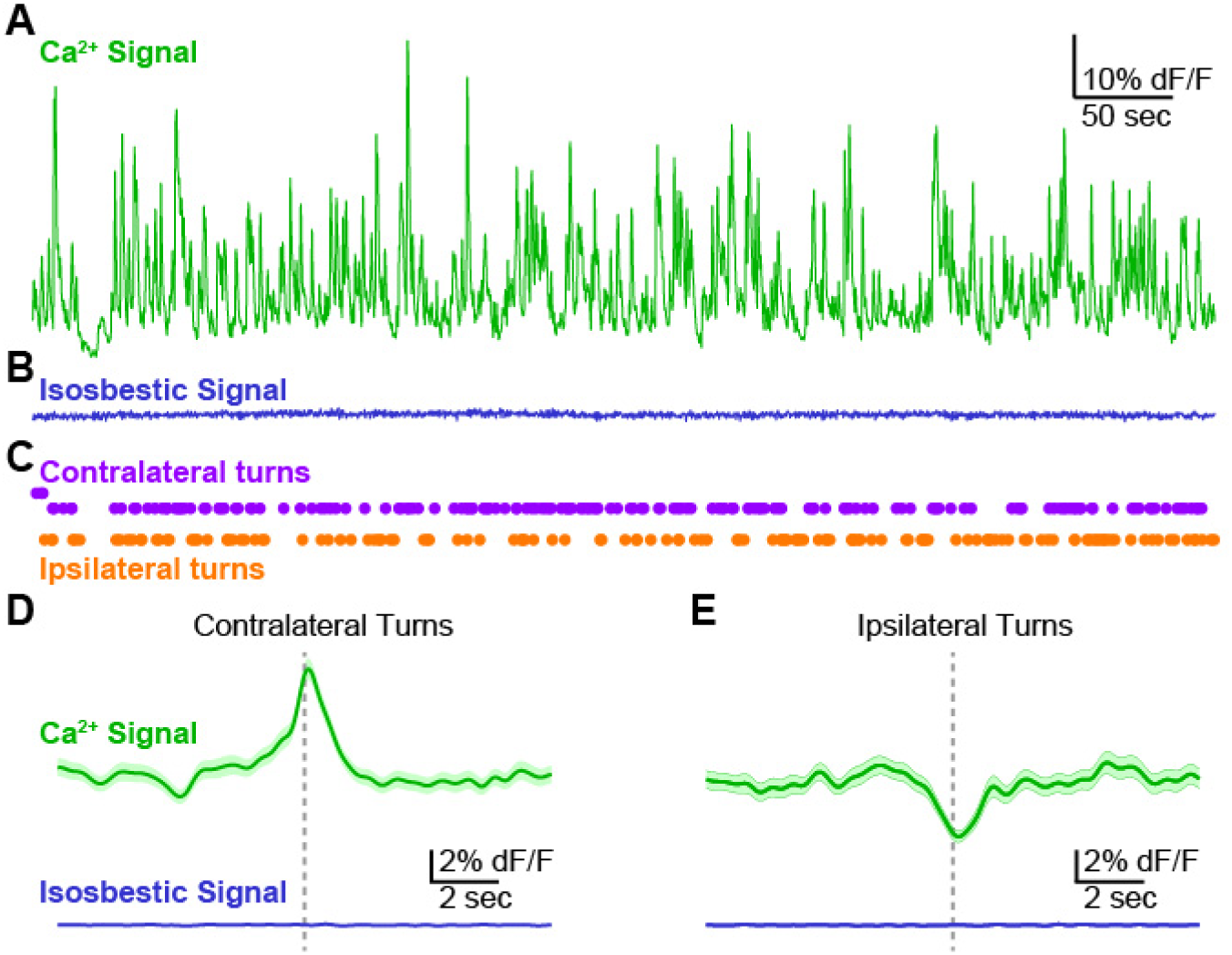
Validation of photometry system by detection of behaviorally aligned calcium signals. A, Exemplar recording (10 min) from dorsomedial striatum of an A2A-cre mouse expressing GCaMP6m in MSNs of the indirect pathway. Calcium-dependent channel. B, Signal from the control, isosbestic channel recorded simultaneously with the calcium-dependent signal in Panel A. C, Contralateral and ipsilateral turns detected by automated video tracking of mouse behavior during the photometry recording session. D,E, Average fluorescence signal aligned to contralateral (Panel D) and ipsilateral (Panel E) turn events. Signal from calcium-dependent (top, green) and isosbestic control channel (bottom, blue). Average signal (dark line) plotted together with standard error (shaded)

## CONCLUDING REMARKS

Photometry is an increasingly important tool with a growing variety of applications in neuroscience. Optical components are readily available for commercial, open-source, and hybrid systems. The electronics that control these optics, however, are either over-engineered and expensive or lack the flexibility and accessibility to be tailored to specific experimental applications. Here we present a cost-effective, open-source system for controlling the optical components in a two-color photometry system, digitizing the resulting signals, and performing post-hoc analysis.

### Post-hoc analysis is essential for proper interpretation of signals

This system uses a microcontroller-based FFT to provide an on-line “preview” of the signal amplitude and structure, similar to those available on commercial systems. Importantly, this “preview” analysis should not be treated as the final measurement of the fluorescence signal. Analysis of the oscillating fluorescence signal requires processing over discrete time windows that are long enough to contain multiple cycles of each oscillation (Figure 4). On-line analysis can draw only on time points that have already occurred, using a time window that extends into the immediate past. This results in a filter-lag that causes the processed signal to trail the true signal by a duration of approximately half the length of the analysis time window (typically 50- 100 ms). In contrast, post-hoc filtering can use an analysis window that extends to both sides of each time point and draws on zero-lag filtering methods that are not possible with on-line filters. This eliminates filter lag as well as any potential computation or signal processing delays, and ensures that the processed signal more accurately reflects the true fluorescence.

### Applications for two-color photometry

Simultaneous measurement of two separate fluorescence channels offers multiple experimental opportunities. In the specific case of GCaMP indicators, calcium-independent control fluorescence levels can be measured from the calcium sensor itself by utilizing the calcium-independent isosbestic point in the emission spectrum (Lerner, Shilyansky et al. 2015). Alternatively, a static indicator in a second fluorescence channel (e.g. tdTomato in the red channel) can provide a control signal to establish the extent to which fluctuations in the primary fluorescence signal (e.g. GCaMP in the green channel) may arise from contamination by environmental light sources, or movement artifacts from the animal, the implant, the optical fiber, or the experimental apparatus. Perhaps most promisingly, however, recent improvements in the latest generation of red-shifted calcium indicators (Zhao, Araki et al. 2011, Akerboom, Carreras Calderon et al. 2013, Wu, Abdelfattah et al. 2014, Inoue, Takeuchi et al. 2015, Dana, Mohar et al. 2016) complement the widely used GCaMP indicators, allowing dynamic interactions between distinct, genetically-targeted neuronal populations to be studied in a single preparation (Markowitz, Gillis et al. 2018).

### A growing toolbox of sensors is expanding applications for one- and two-color photometry

Although GCaMP and other calcium indicators remain the most widely used sensors in photometry experiments (Cui, Jun et al. 2013, Gunaydin, Grosenick et al. 2014, Lerner, Shilyansky et al. 2015, Kim, Yang et al. 2016, Meng, Zhou et al. 2018), emerging genetically-encoded fluorescent sensors for intracellular chloride (Wimmer, Schmitt et al. 2015), voltage (Gong, Huang et al. 2015), and neuromodulators including acetylcholine (Jing, Zhang et al. 2018), dopamine (Patriarchi, Cho et al. 2018, Sun, Zeng et al. 2018) and others are expanding the versatility of this technique by permitting sensitive and specific detection of local neuromodulators through fluorescence signals. The simultaneous development of these new modulatory sensors alongside improved red-shifted calcium indicators (Zhao, Araki et al. 2011, Li and Tsien 2012, Akerboom, Carreras Calderon et al. 2013, Wu, Abdelfattah et al. 2014, Inoue, Takeuchi et al. 2015, Dana, Mohar et al. 2016, Meng, Zhou et al. 2018) suggests it will now be possible to use two-color photometry to simultaneously measure modulatory signaling and its impact on specific neuronal populations with a cellular, spatial and temporal precision that was previously inaccessible in freely moving animals. The low-cost, open-source system described here is designed to complement commercially available optical components to create a flexible system capable of exploiting this new and emerging array of tools for a broad range of neuroscience experiments. Notably, it is also possible to use the same indicator (e.g. GCaMP) in two spatially distinct but nearby regions in the brain and use separate oscillation frequencies of the excitation light in each location to limit fluorescence cross-talk between the two signals.This approach is not possible with a spectral unmixing approach.

### Open-source design supports experimental flexibility

The system described here consists of hardware and firmware for control electronics and processing of signals. We anticipate that these components will provide a valuable complement to the recently published open-source tools that focus primarily on optical and opto-electronic components for similar systems (Simone, Fuzesi et al. 2018). Furthermore, it should be relatively straightforward to integrate both packages into a single open-source design. Although the number of available input and output channels on the microcontroller and the digitizer limit the number of fluorescence channels that can be controlled by a single system using this design, there is no fundamental barrier to using two or more control boxes in parallel within a single experiment if more independent channels of fluorescence recording are required. In this case, the most straightforward solution may be to use two or more of the microcontroller boxes with minimal modifications to each box, and to replace the digitizer with a single 12- or 16-channel digitizer capable of simultaneously acquiring all necessary channels simultaneously. The current design, with all channels accessible through BNC connectors on the front of each box, should facilitate this expansion with minimal additional engineering.

The open-source design of this system allows for unlimited customization to suit specific applications. For example, in long-duration recordings (e.g. over several days for circadian rhythm, automated learning or extended behavior experiments) it may be desirable to include an automated, electronic gating of fluorescent excitation light to minimize photobleaching and phototoxicity. This change can be implemented in minutes by making a simple firmware change to use the “Sync In” BNC connector to gate fluorescence excitation light. Additional changes are possible through modification of the available microcontroller code. For example, a user might implement a firmware system to detect peaks when the fluorescence signal exceeds a certain threshold and have the microcontroller report those peaks with a TTL output for closed-loop control of physiological or behavioral equipment within an experiment. We anticipate that this type of experimental flexibility will be invaluable given the rapidly expanding popularity and range of applications for photometry across neuroscience and other related fields.

## METHODS

### Design and construction of custom-built hardware

The hardware was initially assembled using a prototyping board, short lengths of hook-up wire and direct soldering of individual components. This stage was essential to identify, troubleshoot and correct design flaws. For example, in the first iteration the pulse-width modulated outputs from the MBED microcontroller were connected directly to the analog inputs on the digitizer. This resulted in aliasing artifacts that were eliminated by introducing an RC filter and a voltage follower amplifier on each channel in the final design. Once the design was finalized and validated, a printed circuit board (PCB) was designed and independently tested.

### Microcontroller firmware design

The microcontroller firmware was written using the ARM MBED online compiler (http://os.mbed.com/compiler). The RTOS library (http://os.mbed.com/handbook/RTOS) was imported to support multi-threaded operation. The FFT function was adapted from standard C++ implementations of Fast Fourier Transforms (e.g. http://www.drdobbs.com/cpp). All other functions were written specifically for this application. A pre-compiled binary file is available for download that can simply be copied to the MBED microcontroller by “drag-and-drop” over a USB connection from any modern PC. The code is available here https://hackaday.io/project/160397 for users who wish to customize, modify or improve any aspect.

### Animals

1 adult transgenic mouse on a C57BL/6 background aged 3 months was used in the proof-of-principle open-field recording experiment.

### Stereotactic surgery

All procedures were in accordance with protocols approved by the UCSF Institutional Animal Care and Use Committee. The mouse was maintained on a 12/12 light/dark cycle and fed *ad libitum*. Experiments were carried out during the dark cycle. The surgery was carried out in aseptic conditions while the mouse was anaesthetized with isoflurane (5% for induction, 0.5- 1.5% afterward) in a manual stereotactic frame (Kopf). Buprenorphine HCl (0.1 mg kg^−1^, intraperotineal injection) and Ketoprofen (5 mg kg^−1^, subcutaneous injection) were used for postoperative analgesia. The mouse was allowed to recover for ˜3 weeks before recording.

### Virus injection

We injected 1 µL of adeno-associated virus serotype 1 (AAV1) carrying the calcium indicator GCaMP6m in a double-floxed inverted open reading frame under the control of the Synapsin promoter (AAV1-hSyn-Flex-GCaMP6m). Virus was obtained from the University of Pennsylvania Vector Core. The virus was injected bilaterally into dorsal striatum of an adult mouse in a stereotactic surgery as described above, at coordinates +1.0 anteroposterior (AP),+/-1.5 mediolateral (ML), and −2.5 dorsoventral (DV), measured from bregma on the skull surface. Injections were performed using a glass injector pipette and a Micro-4 Injector system (World Precision Instruments, Inc). The needle was held in place for 1 min before the start of injection, injection speed was 100 nL min^−1^, and the injection needle was raised 5 minutes after completion of virus delivery.

### Optical fiber implants

After virus injection (as described above), a pair of optical fibers (0.48 NA; 400 µm diameter) epoxied to stainless steel fiber optic ferrules (2.5 mm diameter) were implanted in the same surgery. Fiber tip was placed approximately 100 µm above the center of the virus infection zone, at coordinates +1.0 AP, +/-1.5 ML, −2.4 DV from bregma. Dental adhesive (C&B Metabond, Parkell) was used to fix the ferrule in place and coat the surface of the skull. Finally, the skull surface and implant were coated with dental acrylic (Ortho-Jet, Lang Dental). After the cement dried, the scalp was sutured shut.

### Open field behavior tracking

Locomotion was tracked in a brightly lit open-field arena using an overhead camera and post-hoc tracking software (Noldus, Inc). Videos of behavior were recorded from overhead and side views. Rotations (continuous rotational movements over >90 degrees) were tracked using automated detection of the nose and tail positions from post-hoc video analysis.

Synchronization TTL pulses were generated once per minute and digitized by the photometry system alongside the fluorescence measurement to align fluorescence measurements to behavioral events. These pulses were digitized together with the photometry signal, using the “Sync In” input on the photometry system. These pulses were detected off-line with a post-hoc analysis script and used to align the time-stamps from the photometry recording to the time-stamps in the behavioral tracking software (Noldus Ethovision 10). The time of each rotation was then converted to the equivalent time in the photometry recording.

### Fluorescence measurements in vivo

Fluorescence excitation light was generated using two independently driven LEDs. The driver signal for the violet LED (405 nm) was oscillated at 319 Hz to generate excitation light for the isosbestic fluorescence channel. The driver signal for the blue LED (465 nm) was oscillated at 217 Hz to generate excitation light for the calcium-dependent fluorescence channel. Both fiber-coupled LEDs were purchased from Doric Lenses. The photometry control box described in this publication was connected to the LED driver by a pair of BNC cables to deliver the driver signal. The fluorescence mini cube (Doric Lenses) was connected to the intracranial implant on the mouse’s head through an optical fiber (400 µm diameter) and a single channel optical commutator (Doric Lenses). Fluorescence emission light was collected from the intracranial implant through the same optical fiber and detected using a pair of Newport 2151 Femtowatt photodetectors (Newport Inc) connected to the fluorescence mini-cube with optical fibers. The Femtowatt detectors were connected to the photometry box “Photodiode” inputs by BNC connectors for digitization and analysis of the fluorescence signal.

### Signal digitization and off-line analysis

The signal from the two photodiodes was split and sent directly to the digitizer for recording of the raw signal, and to the microcontroller for on-line estimation of the signal amplitude. All signals, including estimated fluorescence (channels 1 and 2), LED driver (channels 3 and 4), raw fluorescence (channels 5 and 6), and TTL synchronization pulses (channels 7 and 8) were digitized continuously at 5 kHz throughout the duration of the experiment. Raw traces plotted in Figure 3A-F were low-pass filtered at 400 Hz for clarity of presentation. In addition to generating the oscillating sine wave driver signals for each fluorescence excitation LED, the microcontroller performs an on-line calculation of the approximate signal amplitudes using a discrete window Fast-Fourier Transform. This signal is valuable when optimizing experimental conditions and assessing the progress of a recording in situ. For final results, however, we strongly recommend off-line post-hoc analysis of the raw fluorescence signals. This allows for better temporal fidelity by using sliding filter windows and removal of filter lags by zero-lag filtering (which is impossible in any on-line filter). A Matlab script (available for download with the rest of the materials) will accomplish this, reading in data directly from a WinEDR data file and processing the results to pull out the signal with a zero-lag filter.

## Data Availability

All data are available upon request. Full parts lists, assembly instructions, compiled firmware, source code and analysis scripts are freely available here: https://hackaday.io/project/160397

## ACKNOWLEDGEMENTS

We thank Benjamin Margolin and Aphroditi Mamaligas for assistance with genotyping and testing, and Mattias Karlsson for technical discussions. Thank you to Alexxai Kravitz, Richard Tsien, Thomas Davidson and the A.C.K. laboratory for comments on the manuscript. This work was funded by NIH R01 NS078435, F32 NS083369 (to S.F.O.) and K99 MH110597 (to S.F.O.),and RR018928 (to the Gladstone Institutes)

## AUTHOR CONTRIBUTIONS

S.F.O. designed the hardware, software and firmware, constructed the apparatus, performed experiments, analyzed data and wrote the manuscript. A.C.K. designed experiments and wrote the manuscript.

## SUPPLEMENTARY INFORMATION

**Table S1.**
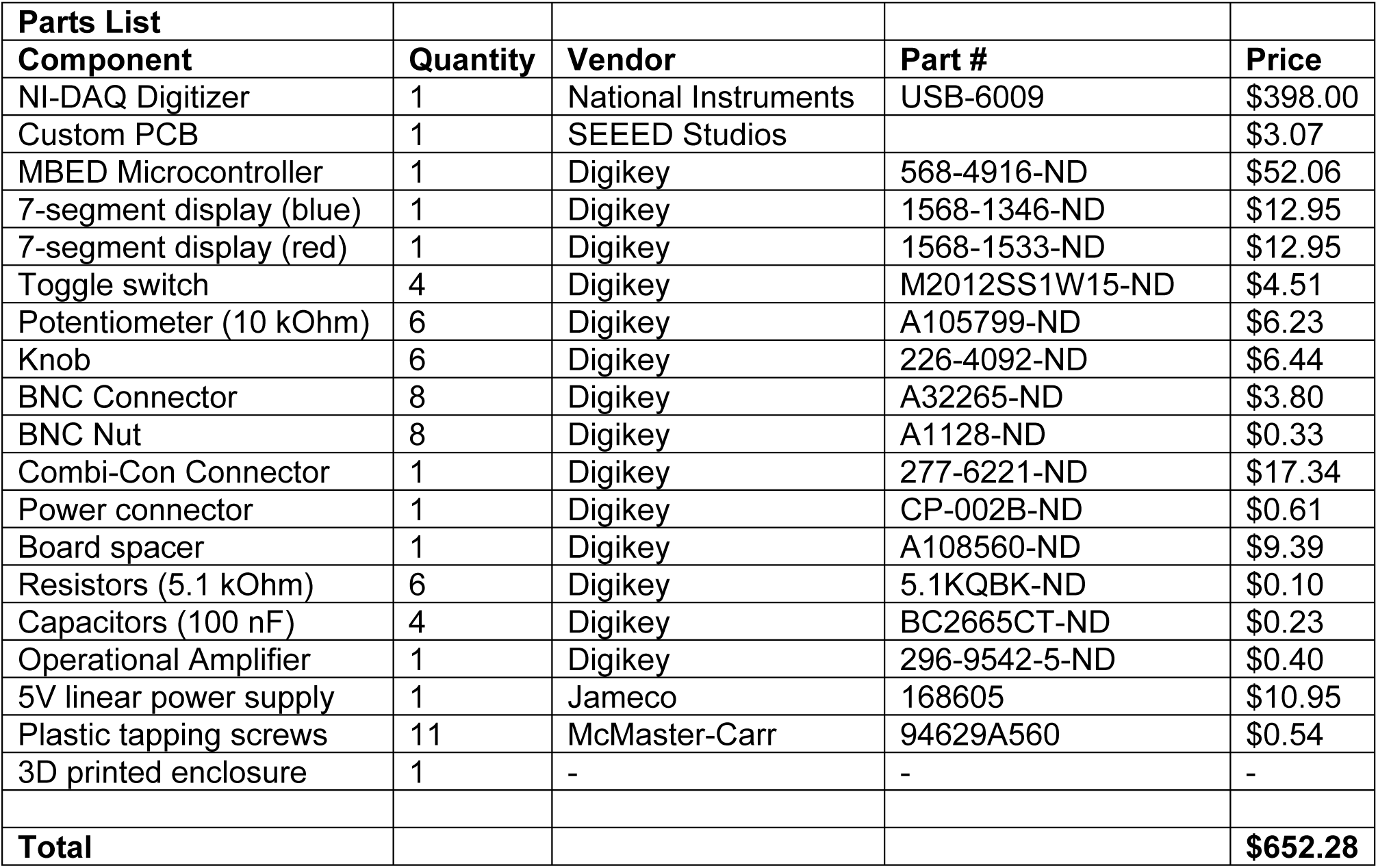
Parts List

